# Geographical distribution of *Anopheles stephensi* in eastern Ethiopia

**DOI:** 10.1101/802587

**Authors:** Meshesha Balkew, Peter Mumba, Dereje Dengela, Gedeon Yohannes, Dejene Getachew, Solomon Yared, Sheleme Chibsa, Matthew Murphy, Kristen George, Karen Lopez, Daniel Janies, Sae Hee Choi, Joseph Spear, Seth R. Irish, Tamar E. Carter

**Author notes:** Corresponding author, Contact: Meshesha Balkew, PMI VectorLink Project, ABC Car Rental Building, Opposite Harmony Hotel, Bole Sub City Woreda 03, Edna Mall, P.O. BOX 13646 Addis Abba Ethiopia.

## Abstract

**Background:** The recent detection of the South Asian malaria vector *An. stephensi* in Ethiopia and other regions in the Horn of Africa has raised concerns about its potential impact on malaria transmission. We report here findings of survey for this species in eastern Ethiopia using both morphological and molecular methods for species identification.

**Methods:** Adult and larval/pupal collections were conducted at ten sites in eastern Ethiopia and *Anopheles* specimens’ species were determined using standard morphological keys and genetic analysis.

**Results:** In total, 2,231 morphologically identified *An. stephensi* were collected. A molecular approach incorporating both PCR endpoint assay and sequencing of portions of the internal transcribed spacer 2 (ITS2) and cytochrome oxidase I (COI) loci confirmed the identity of the *An. stephensi* in most cases (119/124 of the morphologically identified *An. stephensi* confirmed molecularly). Additionally, we observed *Aedes aegypti* larvae and pupae at many of the *An. stephensi* larval habitats.

**Conclusions:** Our findings show that *An. stephensi* is widely distributed in eastern Ethiopia and highlight the need for further surveillance in the southern, western and northern parts of the country and throughout the Horn of Africa.

## BACKGROUND

Malaria remains a leading global health concern with over 250 million cases reported yearly (WHO, 2018). In Ethiopia, even though there has been steady progress in the reduction of malaria (Deribew et al. 2017), 1.5 million cases were reported in 2018 (WHO, 2018). Developing effective malaria control strategies in Ethiopia require knowledge of local mosquito vector species (Animut et al. 2018). One threat to continued progress against malaria is the expansion of vectors into new areas. The South Asian vector *Anopheles stephensi* was recently discovered in Ethiopia and is raising concerns about the impact on malaria transmission in the country and the rest of the Horn of Africa. *Anopheles stephensi* is a major malaria vector of South Asia and the Middle East, including the Arabian Peninsula (Sinka et al. 2011), and is known to transmit both the major malaria parasite species *Plasmodium falciparum* and *P. vivax* (Korgaonkar et al. 2012, Thomas et al. 2017). The first report of *An. stephensi* in the Horn of Africa was from Djibouti in 2013 (Faulde et al. 2014) and was recently confirmed to be persisting in the country (Seyfarth et al. 2019). *An. stephensi* was detected in Ethiopia for the first time in 2016 in Kebridehar (Somali Region) but it remains unclear how broadly distributed the species is in the rest of the country (Carter et al. 2018).

Understanding the distribution of *An. stephensi* in Ethiopia is critical to evaluating the threat it poses to malaria control in Ethiopia and the rest of the Horn of Africa (Carter et al. 2018). It is important during initial surveillance of a potential new vector to evaluate the accuracy of species identifications. Genetic analysis can be a useful complement to morphological identification to achieve optimal accuracy in species identification (Lobo et al. 2015), particularly when identifying a recently detected species. The objective of the study was to investigate the geographic distribution of *An. stephensi* in north eastern and eastern urban localities in Ethiopia using morphological and molecular identification of wild-caught *Anopheles*. With this combined approach, we confirmed that *An. stephensi* is broadly distributed in urban areas in eastern Ethiopia.

## MATERIALS AND METHODS

### Survey sites

*Anopheles stephensi* surveys were conducted from August to November 2018 in ten selected urban sites situated in a climatic zone of either tropical, hot semi-arid or desert with an elevation range of 294 to 2055 meters above sea level. The localities included five in Somali region, three in Afar, one in Amhara region, and Dire Dawa city (Table 1 and Figure 1).

**Table 1.** Collection site altitude and geographic coordinates.

| Site | Region | Altitude (meters above sea level) | Latitude and longitude |
| --- | --- | --- | --- |
| Jigjiga | Somali | 1657 | 9.351765, 42.793937 |
| Erer Gota | Somali | 1090 | 9.556231, 41.384098 |
| Kebridehar | Somali | 532 | 6.738965, 44.277143 |
| Godey | Somali | 294 | 5.949427, 43.553259 |
| Degehabur | Somali | 1065 | 8.223305, 43.558849 |
| Semera | Afar | 431 | 11.794834, 41.008401 |
| Gewane | Afar | 617 | 10.166078, 40.646817 |
| Awash Sebat Kilo | Afar | 916 | 8.989149, 40.164715 |
| Dire Dawa | Dire Dawa | 1178 | 9.596280, 41.854259 |
| Bati | Amhara | 2055 | 11.192694, 40.017657 |

**Fig 1.**
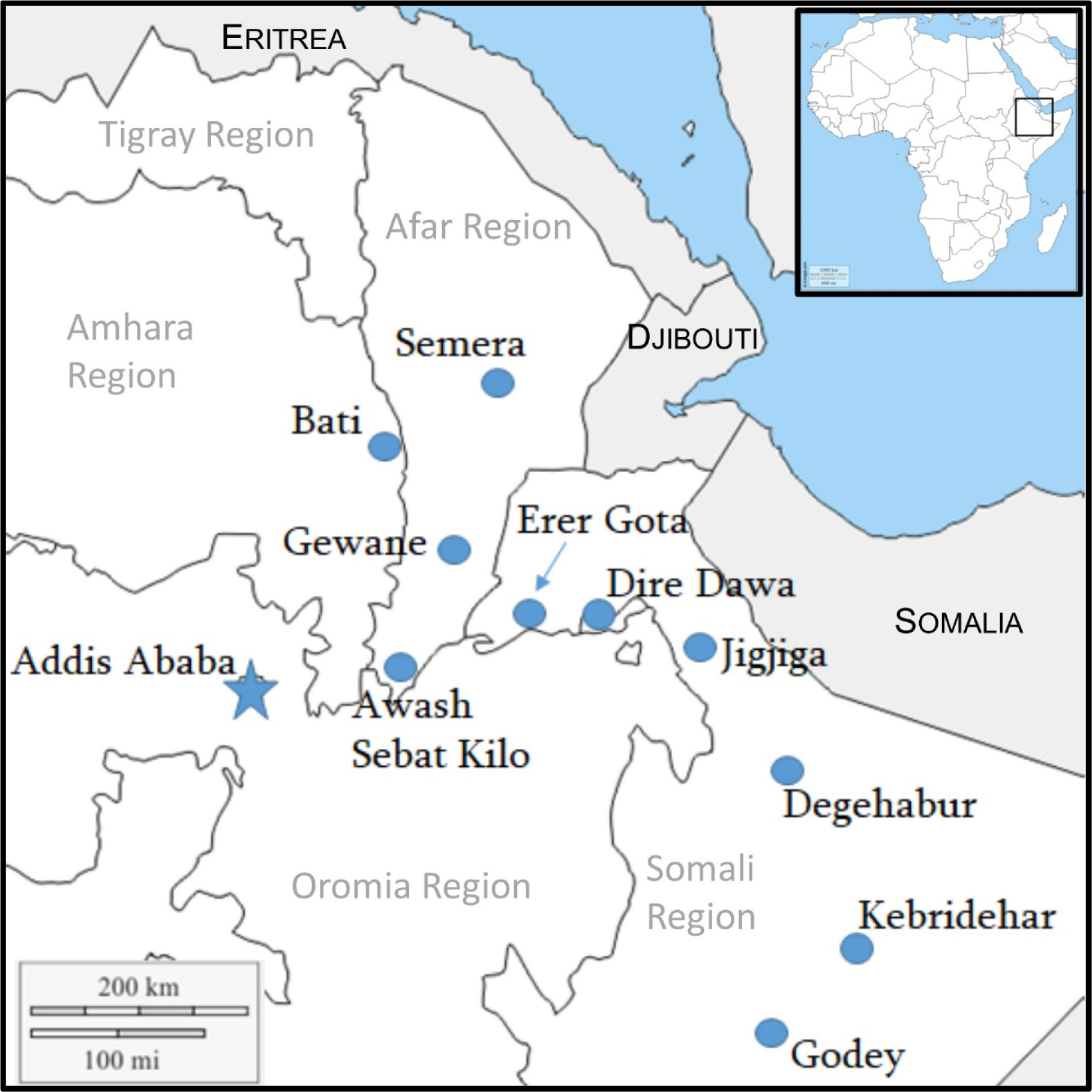
Map of study sites in Ethiopia (blue dots).

**Figure 2.**
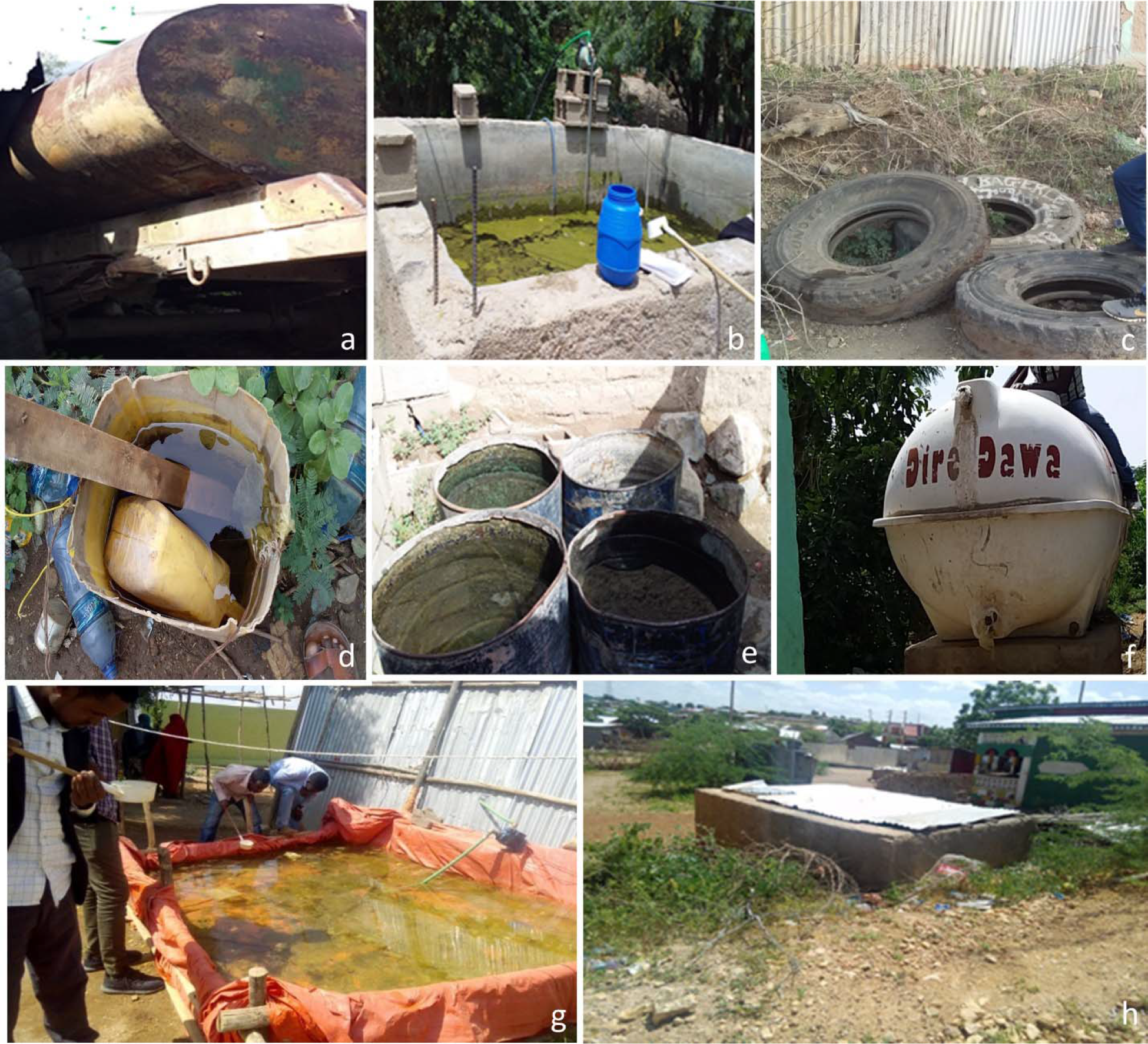
Examples of breeding sites where *An. stephensi* larvae and pupae were found including a) water tank trucks, b) construction water storage reservoirs, c) discarded tires, d) buckets, e) steel drums, f) water tanks, g) temporary water storage reservoirs, h) birkas.

**Table 2.** Number of adult *An. stephensi* collections from PSC, CDC traps, and larval and pupal collections.

| Region | Site | Method of collections and *An. stephensi* |  |  | Total |
| --- | --- | --- | --- | --- | --- |
|  |  | PSC | CDC | Adults Raised from Larvae and Pupae |  |
|  |  | N (%) | N (%) | N (%) |  |
| Somali | Jigjiga | 0 (0) | 0 (0) | 18 (100) | 18 |
|  | Erer | 14 (9.8) | 0 (0) | 129 (90.2) | 143 |
|  | Kebridehar | 3 (0.4) | 0 (0) | 700 (99.6) | 703 |
|  | Godey | 2 (0.6) | 0 (0) | 336 (99.4) | 338 |
|  | Degehabur | 1 (0.6) | 13 (7.2) | 166 (92.2) | 180 |
| Afar | Semera | 38 (12.7) | 1 (0.3) | 260 (87) | 299 |
|  | Gewane | 6 (5.5) | 1 (0.9) | 102 (93.6) | 109 |
|  | Awash Sebat Kilo | 0 (0) | 0 (0) | 26 (100) | 26 |
| Dire Dawa | Dire Dawa | 3 (1.1) | 0 (0) | 277 (98.9) | 280 |
| Amhara | Bati | 0 (0) | 0 (0) | 135 (100) | 135 |
| Total |  | 67 | 15 | 2149 | 2231 |

The areas have mean annual temperatures of about 20°C to 30°C and a mean annual rainfall from 200 to 900 mm. There is a smaller rainy season between March and May, followed by a longer period between July and October (IRI/LDEO Climate Data Library 2019) The sampling was conducted between August and November.

### Sampling of An. stephensi

For the purpose of identification of *An. stephensi* and consequently to determine its presence in the study sites, free flying adult sampling was done together with raising of adults from larval and pupal collections.

### Adult sampling

The entomological methods to sample adult mosquitoes were pyrethrum spray sheet collections (PSC) and Centers for Disease Control (CDC) light traps. In each site, PSC was conducted in 30 houses and CDC light traps were set in 20 houses for one night per house.

Prior to PSC operation, consent was obtained from heads of households and PSC activities were conducted from 6:30 am to 8:00 am mostly in sleeping rooms of big houses and additionally in living rooms of small houses. Houses for PSC were prepared following the WHO standard guidelines. All food and drink was removed, children and small animals made to stay outdoors, and non-removable items together with the floor were covered with pieces of white cloth sheet. All eaves, openings in windows and other mosquito exit points were blocked with pieces of cloth, as much as possible. Two operators, one inside and the other outside of the room/ house were assigned to spray an aerosol insecticide (Baygon, SC Johnson & Son Inc, USA). Wearing a protective nose mask, the operator who was outside the house sprayed the aerosol walking around the house to drive in escapee mosquitoes while the other operator sprayed the insecticide in the whole room by moving from left to right of the door. Then the room/house was closed for ten minutes and knocked down mosquitoes were collected from the ground cloth using forceps and placed in Petri dishes.

CDC light trap collections were made from 6:00 pm to 6:00 am. A CDC light trap was hung 1.5 m above the floor and close to a sleeping location where the occupants are protected with LLINs. Trapped mosquitoes were transferred from collection bag to cages. Those alive were killed with chloroform.

PSC and CDC light trap sampled mosquitoes were sorted into culicines and *Anopheles*. The revised morphological key of Gillies and Coetzee (1987) together with a key prepared by Maureen Coetzee (submitted) were used to discriminate *An. stephensi* from other *Anopheles* particularly *An. arabiensis*. The morphological features that were used to distinguish *An. stephensi* from other *Anopheles* particularly *An. arabiensis* were: speckling on palps, base and apex of palp four with white scales, absence of upper proepisternal setae on the thorax and 3^rd^ main dark area (preapical dark spot) of wing vein R1 with no pale interruption and having two pale spots on the second main dark area of the costa.

Those identified as *An. stephensi* were stored individually in Eppendorf tubes with silica gel to ensure the mosquitoes were kept dry for subsequent molecular analysis.

### Larval and pupal sampling

Larvae and pupae of *Anopheles* were dipped from likely larval breeding habitats including man-made water containers, fresh water pools, stream margins, discarded tires, plastic containers, and irrigation ditches. Water storage for household use and construction is common in the sites. These include metal and plastic tanks, cisterns, barrels and plastic sheets hung from polls. In the Somali region, water is stored in a container locally called “Birka” and it is constructed from cement and stone.

Larvae were reared in field insectaries, feeding them with baking yeast and exposing them to sunlight during the day. Pupae were transferred into adult emergence cages and adults were identified to species using keys described above. Those specimens of *An. stephensi* were preserved as described above.

### Molecular identification of species

To confirm morphological species identifications, a subset of the *An. stephensi* specimens were molecularly characterized. Additional *An. gambiae* s.l. specimens were also analyzed as controls for comparison. Species identification was completed using two approaches: 1) PCR endpoint assay utilizing the internal transcribed spacer 2 (ITS2) locus and 2) sequencing portions of ITS2 and cytochrome oxidase subunit I (COI). ITS2 endpoint assay was performed as previously described (Djadid et al. 2006) using the primers 5.8SB ATGCTTAAATTTAGGGGGTAGTC and 28SC GTCTCGCGACTGCAACTG and the following modifications: Final reagent concentrations and components were 0.5 μM for each primer, 1X Promega GoTAQ HotStart master mix (Promega, Madison, Wisconsin), and water for a total reaction volume of 25ul. The temperature protocol was performed as follows: 95 °C for 1 min, 30 cycles of 96 °C for 30 sec, 48 °C for 30 sec, 72 °C for 1 min, and final extension of 72 °C for 10 min. *Anopheles stephensi* specimens were identified using the visualization of 522 bp band with gel electrophoresis; non-*An. stephensi* specimens will not amplify and no band will be present. Portions of the ITS2 and COI loci were also amplified for sequencing using previously detailed methods (Carter et al. 2019). PCR products were sequenced using Sanger technology with ABI BigDyeTM terminator v3.1 chemistry (Thermofisher, Santa Clara, Ca) according to manufacturer recommendations and run on a 3130 Genetic Analyzer (Thermofisher, Santa Clara, CA). Sequences were cleaned and analyzed using CodonCode (CodonCode Corporation, Centerville, MA). ITS2 and COI sequences from *Anopheles* specimens were submitted as queries to the National Center for Biotechnology Information’s (NCBI) Basic Local Alignment Search Tool (BLAST) (Altschul et al. 1990) against the nucleotide collection in NCBI’s Genbank under default parameters [max High-scoring Segment Pairs (HSP) 250, expect threshold 10, word size 28, optimized for highly similar hits, not specific to any organism]. The *Anopheles* subject sequences from NCBI that formed HSP with the queries were used as the basis of species identification. The percent of species correctly identified using morphology was calculated using the molecular data for comparison.

### *Plasmodium* detection

Wild-caught adult *An. stephensi* were screened for *P. falciparum and P. vivax* DNA as an indication of *Plasmodium* infection. PCR amplification targeted the ssu RNA gene with species specific primers using a previously published approach (Snounou et al. 1993). Presence of a band was positive indication of *Plasmodium* DNA in the sample. *P. falciparum* and *P. vivax* ssu RNA positive controls were included for each reaction set (Microbiologics, St. Cloud, MN, USA).

## RESULTS

A total of 82 adult *An. stephensi* from 300 PSCs and 200 CDC light traps were collected in seven of the 10 sites. The sites with no adult collections were Jigjiga, Awash Sebat Kilo and Bati (Table 1). Of the 82 adults 81.7% (N=67) were from PSC and the remaining 18.3% (N=15) were from CDC light traps. The majority of *An. stephensi* sampled using PSC were from Semera and Erer and that of CDC were from Degehabur. Larval and pupal collections yielded 2,149 adult *An. stephensi* from all the sites confirming the presence of immature stages (Table 1). In addition to *An. stephensi*, *Aedes aegypti* larvae and pupae were detected.

PCR endpoint assay was performed, and successful PCR products were obtained for 130/133 attempted *Anopheles.* With the PCR endpoint assay, 119 specimens were identified as *An. stephensi* and 11 specimens were identified as *non-An. stephensi*. Sequencing of portions of the ITS2 and cytochrome oxidase subunit I (COI) loci was also completed and successful sequencing was completed for 118 *Anopheles*. BLAST analysis of *Anopheles* sequences confirmed the positive detection of *An. stephensi* at all ten sites. Sequence-based species identification was mostly consistent with ITS2 endpoint assay results, with the exception of a single specimen that was morphologically identified as *An. gambiae* s.l., identified as non-*An. stephensi* with endpoint assay, but sequencing detected *An. stephensi*. BLAST of ITS2 sequences further identified all the non-*An. stephensi* specimens as *An. arabiensis*.

We compared the morphological identification to the ITS2 PCR endpoint assay results. Of the 130 *Anopheles* included, 124 were classified as *An. stephensi* and six as *An. gambiae* s.l. based on morphology. Five of the 124 (4.0%) morphologically identified *An. stephensi* were not confirmed to be *An. stephensi* with the PCR endpoint-assay. All morphologically identified *An. gambiae* s.l. (sequence-confirmed *An. arabiensis*) that were successfully amplified were also identified as non-*An. stephensi* with the ITS2 PCR endpoint assay.

### *Plasmodium* detection

All 82 wild-caught adult *An. stephensi* were screened for *P. falciparum or P. vivax* infection. No *Plasmodium* DNA was detected in any specimens.

## DISCUSSION

This survey confirms that *An. stephensi* is distributed broadly in eastern Ethiopia. These data, taken with previous reports of *An. stephensi* in Kebridehar in 2016 (Carter, Yared et al. 2018), confirm that *An. stephensi* is established in this region. This is the first evidence for the presence of adult *An. stephensi* multiple regions in Ethiopia, where it might transmit malaria. The broad distribution in Ethiopia as far south as Godey (about 600 km south of Djibouti city) suggests that if *An. stephensi* is a relatively new introduction into Ethiopia, it has spread quickly. Alternatively, it may have been in Ethiopia for years before its detection in 2016. The widespread presence of *An. stephensi* in Ethiopia along with Djibouti suggests that neighboring countries, such as Sudan, South Sudan, Eritrea, Somalia, and Kenya, should also enhance surveillance. Given that *An. stephensi* in Djibouti was found to carry both *P. falciparum* and *P. vivax*, there is potential for the same to be observed in Ethiopia, thus malaria control strategies should now consider the potential for the established *An. stephensi* to transmit malaria.

The presence of *An. stephensi* was confirmed both morphologically and molecularly. While morphology was mostly consistent with molecular approach (119/124 correctly identified *An. stephensi*), there were a few instances of incorrectly identified morphologies highlighting the risk for the misidentification of specimens. As more vector surveillance programs in Africa incorporate *An. stephensi* into their morphological keys, molecular data can be helpful with the evaluation of the successful training in *An. stephensi* morphological identification. This is particularly important at this beginning phase of *An. stephensi* surveillance as field technicians will be adapting to detecting *An. stephensi*. We incorporated two molecular approaches, one of which, the end-point assay using primers designed by Djadid et al (2006) is more feasible in resource limited settings. We found that this assay was mostly consistent with the sequence data and has to potential be integrated into current PCR based assays that focus on the detection of members of *An. gambiae* complex, the most common malaria vectors in Africa.

While we have confirmed the broad distribution of *An. stephensi* in eastern Ethiopia, the distribution in the western part of the country has yet to be determined. West Ethiopia has had more consistent surveillance of malaria vectors than the eastern, due to the burden of the disease there, however previously used trapping methods may limit the ability to detect *An. stephensi*. The current trapping techniques that rely heavily on CDC light traps may limit the ability to detect *An. stephensi*, given the low number of *An. stephensi* caught with CDC light traps at most sites in this study. Additional studies on the breeding, feeding, and resting behavior of *An. stephensi* can provide crucial information that can be applied to enhance future surveillance efforts in west and eastern Ethiopia.

Several additional areas of query need to be pursued further to better inform vector control efforts. No study has been published to confirm that the Ethiopian *An. stephensi* can or does transmit *Plasmodium*. Both field confirmation of infected *An. stephensi* and laboratory infection are helpful approaches evaluating this information. In this study, the 82 wild-caught *An. stephensi* in this study were screened for both *P. falciparum* and *P. vivax* using PCR and *Plasmodium* was not detected. This is not unexpected as the regions included in this study report low malaria transmission, so a much larger sample size would be required to detect *Plasmodium* infection in *An. stephensi*. Future surveillance will continue to screen for *Plasmodium* using both a PCR-based and circumsporozoite protein Enzyme Linked Immunosorbent Assay (ELISA).

The surveillance of *An. stephensi* to this point has been conducted in short time spans, with limited ability to assess changes in *An. stephensi* population size over time. We will be repeating collections in multiple sites in eastern Ethiopia to provide crucial information about how the population is changing year to year. This information will be particularly important as new vector control interventions are rolled out to evaluate their effectiveness. Insecticide resistance has been reported in the dominant malaria vector *An. arabiensis* in Ethiopia (Messenger et al. 2017), but insecticide resistance status in *An. stephensi* is unknown. Investigations into insecticide resistance, the molecular mechanisms behind resistance, and potential genetic markers that can be used for surveillance are ongoing.

One lingering question relates to the origin of *An. stephensi* in Ethiopia. Previous phylogeographic analysis revealed the closest COI sequence similarity of the Ethiopia *An. stephensi* found in Kebridehar to a specimen from Pakistan (Carter et al. 2018). Phylogeographic analysis including sequencing from recent global *An. stephensi* collections using multiple loci or whole genome sequences can help identify the exact origin of the *An. stephensi* in the Horn of Africa and how it has spread throughout the region. This information will support efforts to prevent the further introduction and spread of *An. stephensi*.

An ancillary observation during the surveillance was the detection of larvae of the dengue vector *Aedes aegypti* together with *An. stephensi* larvae, suggesting that these two vectors share larval habitats. Dengue is a growing public health concern in Ethiopia, particularly in eastern Ethiopia, where major outbreaks were reported in eastern Ethiopia in 2013 (Woyessa et al. 2014, Yusuf et al. 2016) and 2015 (Degife et al. 2019). With the finding of *Ae. aegypti* larvae with *An. stephensi*, we can consider integrated vector control to target both *An. stephensi* and *Ae. aegypti.* This would be a cost-effective approach to reducing both malaria and dengue virus transmission. Future surveillance in eastern Ethiopia will work towards determining relative abundance of *Ae. aegypti* larvae at *An. stephensi* breeding sites.

## CONCLUSION

We confirmed that *An. stephensi* is widely distributed and established in eastern Ethiopia. Studies are ongoing to evaluate the distribution in the rest of the country and the potential risk for *An. stephensi* to change the malaria transmission landscape in the country and the rest of the African continent. Cross country cooperation and collaborations are needed to effectively address this potential global health concern.

## LIST OF ABBREVIATIONS

BLAST: Basic Local Alignment Search Tool
CDC: Centers for Disease Control and Prevention
CO1: Cytochrome Oxidase Subunit 1
DNA: deoxyribonucleic acid
ELISA: Enzyme Linked Immunosorbent Assay
HSP: High-scoring Segment Pairs
ITS2: internal transcribed spacer 2
NCBI: National Center for Biotechnology Information
PCR: Polymerase Chain Reaction
PMI: U.S. President’s Malaria Initiative
PSC: Pyrethrum spray catch
RNA: Ribonucleic acid
ssu: small subunit
USAID: United States Agency of International Development
WHO: World Health Organization

## Ethics approval and consent to participate

Wild mosquitoes used for this study were collected from dwellings and animal houses, following homeowners’ verbal consent. The study protocol was reviewed by the Centers for Disease Control and Prevention, USA and deemed to be non-human research (2017-227).

## Consent for publication

Not applicable

## Availability of data and materials

All data generated or analysed during this study are available from the corresponding author upon request.

## Authors’ contributions

MB, PM, DD, SC, MM, KG, SI, TC developed the conception and design of the study. MB, GY, DG, SY collected the field data and identified mosquito samples. KL, DJ, SHC, JS, and TC conducted and interpreted the laboratory analysis. MB and TC analyzed the data. MB, SI, and TC contributed to the writing of the paper. All authors read and approved the final manuscript.

## Acknowledgements

Cecilia Flately (Technical Program Manager, Abt Associates,, PMI VectorLink Project, Rockville, MD) and Getinet Awoke (Environmental Compliance Officer, Abt Associates, PMI VectorLink Project, Addis Ababa, Ethiopia) are thanked for their assistance to the project.

## REFERENCES

Ahmed YM, Salah AA. Epidemiology of dengue fever in Ethiopian Somali region: retrospective health facility based study. Cent Afr J Public Health. 2016; 2:51–6.

Altschul SF, Gish W, Miller W, Myers EW, Lipman DJ. Basic local alignment search tool. J Mol Biol. 1990; 215:403–10.

Animut A, Lindtjørn B. Use of epidemiological and entomological tools in the control and elimination of malaria in Ethiopia. Malar J. 2018; 17:26.

Carter TE, Yared S, Gebresilassie A, Bonnell V, Damodaran L, Lopez K, et al. First detection of *Anopheles stephensi* Liston, 1901 (Diptera: Culicidae) in Ethiopia using molecular and morphological approaches. Acta Trop. 2018; 188:180–6.

Carter TE, Yared S, Hansel S, Lopez K, Janies D. Sequence-based identification of *Anopheles* species in eastern Ethiopia. Malar J. 2019; 18:135.

Deribew A, Dejene T, Kebede B, Tessema GA, Melaku YA, Misganaw A, et al. Incidence, prevalence and mortality rates of malaria in Ethiopia from 1990 to 2015: analysis of the global burden of diseases 2015. Malar J. 2017; 16:271.

Faulde MK, Rueda LM, Khaireh BA. First record of the Asian malaria vector *Anopheles stephensi* and its possible role in the resurgence of malaria in Djibouti, Horn of Africa. Acta Trop. 2014; 139:39–43.

Gillies MT, Coetzee M. A Supplement to the Anophelinae of Africa South of the Sahara (Afrotropical Region). Publications of the South African Institute for Medical Research, Johannesburg, no. 55; 1987.

Gillies MT, de Meillon B. The Anophelinae of Africa South of the Sahara (Ethiopian Zoogeographical Region). Publications of the South African Institute for Medical Research, Johannesburg, no. 54; 1968.

Degife LH, Worku Y, Belay D, Bekele A, Hailemariam Z. Factors associated with dengue fever outbreak in Dire Dawa administration city, October, 2015, Ethiopia-case control study. BMC Public Health. 2019; 19:650.

Djadid ND, Gholizadeh S, Aghajari M, Zehi AH, Raeisi A, Zakeri S. Genetic analysis of rDNA-ITS2 and RAPD loci in field populations of the malaria vector, *Anopheles stephensi* (Diptera: Culicidae): implications for the control program in Iran. Acta Trop. 2006; 97:65–74.

IRI/LDEO Climate Data Library. Palisades, NY: Lamont-Doherty Earth Observatory (LDEO) Climate Group, Columbia University. https://iridl.ldeo.columbia.edu/maproom/Global/Climatologies/Select_a_Point.html. Accessed 1 Sept 2019.

Korgaonkar NS, Kumar A, Yadav RS, Kabadi D, Dash AP. Mosquito biting activity on humans & detection of *Plasmodium falciparum* infection in *Anopheles stephensi* in Goa, India. Indian J Med Res. 2012; 135:120.

Lobo NF, St Laurent B, Sikaala CH, Hamainza B, Chanda J, Chinula D, et al. Unexpected diversity of *Anopheles* species in Eastern Zambia: implications for evaluating vector behavior and interventions using molecular tools. Sci Rep. 2015; 5:17952.

Messenger LA, Shililu J, Irish SR, Anshebo GY, Tesfaye AG, Ye-Ebiyo Y, et al. 2017. Insecticide resistance in *Anopheles arabiensis* from Ethiopia (2012-2016): a nationwide study for insecticide resistance monitoring. Malar J 16: 469.

Seyfarth M, Khaireh BA, Abdi AA, Bouh SM, Faulde MK. Five years following first detection of *Anopheles stephensi* (Diptera: Culicidae) in Djibouti, Horn of Africa: populations established—malaria emerging. Parasitol Res. 2019; 118:725–32.

Sinka ME, Bangs MJ, Manguin S, Chareonviriyaphap T, Patil AP, Temperley WH, et al. The dominant *Anopheles* vectors of human malaria in the Asia-Pacific region: occurrence data, distribution maps and bionomic précis. Parasit Vectors. 2011; 4:89.

Snounou G, Viriyakosol S, Jarra W, Thaithong S, Brown KN. Identification of the four human malaria parasite species in field samples by the polymerase chain reaction and detection of a high prevalence of mixed infections. Mol Biochem Parasitol. 1993. 58: 283–292.

Thomas S, Ravishankaran S, Justin NJ, Asokan A, Mathai MT, Valecha N, et al. Resting and feeding preferences of *Anopheles stephensi* in an urban setting, perennial for malaria. Malar J. 2017; 16:111.

World Malaria Report 2018. Geneva: World Health Organization; 2018

Woyessa AB, Mengesha M, Kassa W, Kifle E, Wondabeku M, Girmay A, Kebede A, Jima D. The first acute febrile illness investigation associated with dengue fever in Ethiopia, 2013: a descriptive analysis. Ethiop J Health Dev. 2014; 28:155–61.

